# Discrete depolarisations of chordotonal organ neurons propagate toward the soma and are independent of the sensory TRP channels NompC and Nanchung-Inactive

**DOI:** 10.64898/2026.07.06.735281

**Authors:** Atitheb Chaiyasitdhi, Hao Li, Mingxuan Zhao, Huimin Jing, Quig Wei, Tingting Zhang, Ben Warren

**Affiliations:** Shanxi Key Laboratory of Nucleic Acid Biopesticides, Institute of Applied Biology, Shanxi University, Taiyuan, 030006, China; Department of Biology, Faculty of Science, Mahidol University, Bangkok, Thailand; Center of Exellence for Vectors and Vectors-Borne Diseases, Faculty of Science, Mahidol University, Bangkok, Thailand; Department of Genetics, Genomics, and Cancer Sciences, School of Biological and Biomedical Sciences, University of Leicester, United Kingdom; School of Life Sciences, Keele University, Newcastle, ST5 5BG, United Kingdom

## Abstract

The electrophysiological process of auditory transduction in insects remains largely conjecture due to the unknown role of ion channels localised to the cilia, but experimental evidence supports either NompC or Nan-Iav as the auditory mechanotransduction ion channel. Here, we knocked down two key genes that code for the two candidate sound-activated ion channels using dsRNA-mediated RNA interference. We measured sound-evoked activity of the auditory nerve and intracellular electrical currents from the ciliated ending of individual auditory receptors. We found that the sound-evoked nerve activity was reduced in *nompC, nan* and *ift88* knockdown. Using whole-cell patch clamp recordings we found that *nompC* and *nan* knockdown resulted in reduced sound-evoked transduction current. Stochastic depolarisations hypothesised to be mediated from one of the candidate mechanotransduction ion channels, either NompC or Nan-Iav, where not affected by knockdown of either channel. The discrete depolarisations are therefore mediated through another unidentified ion channel. We test the hypothesis that discrete depolarisations are graded action potentials that travel toward the soma through noise analysis of the transduction current and analysis of discrete depolarisations to voltage-steps. As a positive control we also knocked down *ift88*, a protein essential for transporting proteins, including ion channels, along the cilium and found both the transduction current and the discrete depolarisations decreased.

**Key points:**

- Injection of dsRNA decreased RNA of *nompC* and *nan*
- Sound-evoked nerve activity is reduced for RNAi-mediated knockdown of *nompC* and *nan*
- NompC and Nan both contribute to the transduction current
- The stochastic discrete depolarisations are not due to NompC or Nan-Iav ion channel but to a third unidentified ion channel.
- Noise analysis of the transduction current and the discrete depolarisations suggests they are graded action potentials that travel in the direction of the soma.
- Knockdown of *ift88* reduced both the transduction current and discrete depolarisations.

**Significance Statement:** Insects are important to understand, economically, agriculturally and medically. However, we still do not understand fundamental aspects of how insects detect their own body movements, vibrations and sound. These senses are detected by insect chordotonal organs, specialised miniaturised mechanoreceptors that convert movements into electrical signals through specialised ion channels. Previous experimental work has advocated either NompC or Nan-Iav as the mechanosensitive ion channel. Here, for the first time, we reduced the expression of both *nompC* and *nan* and measured the sound-evoked transduction current directly from neurons in a specialised auditory chordotonal organ. In contradiction to previous studies, we show that both ion channels contribute to the transduction current and find that electrical signals termed “discrete depolarisations” travel toward the soma.

## Introduction

Over the last 25 years the mammalian auditory transduction field has transitioned from the belief of one mechano-electrical transduction (MET) ion channel to the realisation that several ion channels and proteins coded by several genes compose a functioning MET complex (Zheng & Holt, 2021). In insects, however, two contending ion channels are vying to be *the* auditory MET ion channel, two transient receptor potential channels, No mechanoreceptor potential C (NompC) and Nanchung-Inactive (Nan-Iav) (a single ion channel composed of two genes, *nan* and *iav*). These ion channels have been localised in *Drosophila*’s auditory Johnston’s organ, NompC at the tip of the cilium and Nan-Iav in the basal segment suggesting distinct physiological functions (Gong et al., 2004; Lee et al., 2010; Liang et al .,2011). All three genes are expressed in the auditory system of the desert locust (French and Warren, 2021) and insecticides that bind to Nan-Iav, disrupt chordotonal organ function in a variety of insects (Fedor et al., 2026; Wang et al., 2019; Song et al., 2023). This suggests that the genes have similar roles across insect orders.

Early studies of MET relied on mechanical measurements on the antennal sound-receiver of *Drosophila* and extracellular electrical recordings of the auditory nerve. Remarkably properties of the MET ion channel can be deduced from mechanical properties of the antennal sound-receiver to which they attach (Efferzt et al., 2011). This is due to the coordinated opening and closing of thousands of MET ion channels from hundreds of auditory neurons that bestow forces back onto the antenna in a process called active amplification (Göpfert et al., 2005). Genetic knockout of *nompC* leads to an abolition of active amplification and a reduction in sound-evoked nerve activity (Gopfert and Robert, 2003; Gopfert et al., 2006). Genetic knockdown of *nan* or *iav* leads to overactive mechanical amplification and abolished sound-evoked nerve activity (Gopfert et al., 2006). This led to the hypothesis that Nan-Iav (localised along the basal length of the cilium) carries an electrical signal initiated from NompC (at the tip of the cilium) and that Nan-Iav negatively feedbacks to regulate mechanical amplification (Gopfert et al., 2006). Indeed, dynein is widely thought to provide the force for active amplification working similar to known motile cilia and is co-localised in the cilium with Nan-Iav (Eberl et al., 2000; Gong et al., 2004; Warren et al., 2011; Karak et al., 2015; Mockel et al., 2012).

Important properties of both auditory MET channel candidates were determined from expression in heterologous cells or ectopic expression in olfactory-processing cells. NompC forms compression-sensitive ion channel (Yan et al., 2013; Wang et al., 2021), whereas Nan-Iav were not mechanically sensitive (Gong et al., 2004); thus supporting NompC as a bone fide mechanosensitive ion channel. Although Nan-Iav is not mechanically sensitive when heterologously expressed it could still be mechanically sensitive *in vivo*.

Recordings of the transduction current, combined with genetic and pharmacological manipulations of the channel, are required to determine the identity of the insect auditory MET channel. The first such recordings were pioneered in *Drosophila* from recordings of the giant fibre neuron onto which the hundreds of auditory neurons form electrical synapses (Lehnert et al., 2013). Here, knockout of Nan-Iav but not NompC resulted in abolition of sound-evoked summated currents thought to represent the transduction current. Intracellular patch-clamp recordings from auditory neurons were pioneered in the desert locust’s Müller’s organ but these were not combined with genetic manipulation of ion channel expression (Warren & Matheson, 2018). Here, the maximum Nan-Iav current matched the maximum sound-evoked transduction current supporting Nan-Iav as the MET channel. Transduction currents have been recorded from *nompC, nan and iav* mutants in proprioceptive (lch) CO neurons of *Drosophila* larvae (Li et al., 2021). Only knockdown of *nan* or *iav* resulted in decreased transduction current. All four types of CO neurons so far recorded have discrete depolarisations, temporally random depolarisations of varying magnitude (Hill, 1983; Warren & Matheson, 2018; Zhang et al., 2013; Atitheb et al., in preparation). They have been assumed to be stochastic openings of the transduction channels themselves (Warren & Matheson, 2018; Austin et al., 2023) but this has not been explicitly tested.

Here, we combined knockdown of key ion channel genes, *nompC* and *nan* in *Locusta migratoria* with patch-clamp recordings of the sound-evoked transduction current and discrete depolarisations from individual auditory neurons of Müller’s organ *ex vivo*. In the same knockdown locusts, we recorded spontaneous and sound-evoked spike responses from the auditory nerve *in vivo*. We complemented this approach with intracellular recordings from auditory neurons of Müller’s organ in *Schistocerca gregaria* to analyse the noise of the transduction current and magnitude of the discrete depolarisations during a voltage-step.

## Methods

### Locust Husbandry

Migratory locusts, *Locusta migratoria* were obtained from the Kunming Locust Breeding Center were reared in crowded conditions (phase gregaria) on 14 h:10 h light:dark cycle at (30±2)°C and 50% relative humidity. The locusts were provided with a diet of fresh wheat seedlings and wheat bran.

Desert locusts, *Schistocerca gregaria*, from a long-term culture at the University of Leicester were reared in crowded conditions (phase gregaria) on a 12 h light/dark cycle at 36°C and 25°C, respectively. Locusts were fed on a combination of fresh wheat and bran. Mixed sex locusts between 10 and 20 d after imaginal moult were used for all experiments.

### Genetic knockdown and quantification of gene expression

For the knockdown experiments, adult locusts on the first day after eclosion were anesthetized on ice. Approximately 13 µg of dsRNA (dissolved in nuclease-free water) was injected into the hemocoel between the second and third abdominal segments using a micro-syringe. A second injection of 10–12 µg dsRNA was administered on the 7th day after eclosion for boosting RNAi efficiency. Control groups were injected with an equal volume of GFP dsRNA in nuclease-free water for each injection.

To evaluate gene expression levels, we performed reverse transcription quantitative polymerase chain reaction (RT-qPCR) to measure the transcript levels of *nompC, nan* and *ift88*. Müller’s organs were extracted by piercing the external tympanum with fine forceps either side of the folded body. The forceps were closed onto Müller’s organ before it was pulled out, and wiped onto a pestle in an Eppendorf tube submerged in liquid nirotrgen. A total of 12 Müller’s organs from 6 locusts were used for each qPCR reaction. Total RNA was isolated from Müller’s organs on days 2, 3, 6, 8 and 9 days after last moult using RNAiso Plus Kit (TaKaRa, Japan; Cat. #9109) following the manufacturer’s protocol. First-strand cDNA synthesis was conducted with HiScript III RT SuperMix (+gDNA wiper; Vazyme, China; Cat. #R323–01). The cDNA was diluted 5-fold by mixing 20 μL of the sample with 80 μL of sterile, nuclease-free water to RT-qPCR analysis. RT-qPCR was performed using the SYBR Green Real-time PCR Master Mix (Roche, Basel, Switzerland) and Quantitative real-time PCR was performed on a CFX96 Touch™ Real-Time PCR Detection System (Bio-Rad, Hercules, CA, USA). The thermal cycling conditions were as follows: initial denaturation at 95°C for 30s, followed by 40 cycles of denaturation at 95.0°C for 5s, and annealing/extension at 60°C for 30s. A melt curve analysis was generated by heating from 65.0°C to 95.0°C in 0.5°C increments (5s per step) to verify amplicon specificity. The relative expression levels were calculated using the 2^−ΔCt^ method, with *Rpl32* gene serving as the internal reference. All the RT-qPCR reactions were performed in three biological replicates.

### Acoustic stimulation

Sound Pressure Levels (SPLs) were measured with a microphone (Pre-03 Audiomatica, DBS Audio) and amplifier (Clio Pre 01 preamp, DBS Audio). The microphone was calibrated with a B&K Sound Level Calibrator (CAL73, Mouser Electronics). For hook electrode and patch-clamp recordings, the locust ear was stimulated with the same speaker and amplifier as above with a 3 kHz pure tone duration of 0.25s. For hook electrode recordings the 3 kHz tone had a rise and fall time of 2 ms. Tones were played three times for each locust at each SPL and the average response taken for each SPL. For intracellular recordings from individual auditory neurons the speaker was driven by a custom-made amplifier controlled by an EPC10-USB patch-clamp amplifier (HEKA-Elektronik) controlled by the program Patchmaster (version 2×90.2, HEKA-Elektronik) running under Microsoft Windows (version 10).

### In vivo hook electrode recordings from auditory nerve six

Locusts were secured ventral side up in plasticine. The second and third ventral thoracic sternites were surgically removed to expose the metathoracic ganglion, which lies beneath the tracheal air sacs. The tracheae were then carefully removed using fine scissors and forceps. Hook electrodes made from 50 μm diameter copper wire was hooked to the auditory nerve N6, and the nerve was lifted out of the haemolymph. To prevent the nerve from drying out, a mixture of 70% Vaseline and 30% cedar oil was applied to the auditory nerve using a syringe. Neural signals were amplified 1,000× using a differential amplifier (DAM50, World Precision Instruments), filtered with a 300 Hz high-pass filter and a 10 kHz low-pass filter, and sampled at 100 kHz using a National Instruments data acquisition device (NI USB-6221). Data acquisition and acoustic stimulation were controlled using a custom MATLAB GUI-based program running on Windows 11 (MATLAB R2023–2025, MathWorks). Sound-evoked neural responses were quantified by calculating the ratio of the root mean square (RMS) amplitude during the 250 ms sound stimulus to the RMS amplitude of the background neural activity measured during the 50 ms periods immediately before and after stimulus presentation.

### Dissection of Müller’s organ and isolation of Group-III auditory neurons

Whole cell patch clamp recordings were performed on group-III auditory neurons because they form the majority of auditory neurons of Müller’s organ (∼46 out of ∼80) (Jacobs *et al*., 1999), they are the most sensitive auditory neurons of Müller’s organ (Römer, 1976) and are broadly tuned to the 3 kHz we used for noise-exposure (Warren and Matheson, 2018). For intracellular patch-clamp recordings from individual auditory neurons the abdominal ear, including Müller’s Organ attached to the internal side of the tympanum, was excised from the first abdominal segment, by cutting around the small rim of cuticle surrounding the tympanum with a fine razor blade. For *L. migratoria* the “pinnae” flap that partially obscures the auditory cavity was cut at its base and removed. Trachea and the auditory nerve (Nerve 6) were cut with fine scissors (5200-00, Fine Science Tools), and the trachea and connective tissue removed with fine forceps. This preparation allowed perfusion of saline to the internal side of the tympanum, necessary for water-immersion optics for visualizing Müller’s Organ and the auditory neurons to be patch-clamped, and concurrent acoustic stimulation to the dry external side of the tympanum. Dissection, protease and recordings took ∼60 min for each locust ear.

To expose Group-III auditory neurons for patch-clamp recordings, a solution of collagenase (10 mg/mL) and hyaluronidase (10 mg/mL) (C5138, H2126, Sigma Aldrich) in extracellular saline was applied onto the medial-dorsal border of Müller’s organ through a wide (12 μm) patch pipette to digest a small opening in the capsule enclosing Müller’s organ. The patch-electrode was then inserted through the capsule opening to access neuronal somate. For *S. gregaria* the concentration of hyaluronidase was 0.5 mg/ml as the capsule that surrounds neuronal somata in *S. gregaria* is easier to digest. For *S. gregaria* large sections of the capsule was removed to expose the membrane of Group-III auditory neurons. The somata were visualized with a Cerna mini microscope (SFM2, Thor Labs), equipped with infrared LED light source and a water immersion objective (NIR Apo, 40×, 0.8 numerical aperture, 3.5 mm working distance, Nikon) and multiple other custom modifications.

### Electrophysiological recordings and isolation of the transduction current

Electrodes with tip resistances between 3 and 4 MΩ were fashioned from borosilicate glass (0.86 mm inner diameter, 1.5 mm outer diameter; GB150-8P, Science Products GmbH) with a vertical pipette puller (PC-100, Narishige). Recording pipettes were filled with intracellular saline containing the following (in mM): 170 K-aspartate, 4 NaCl, 2 MgCl2, 1 CaCl2, 10 HEPES, 10 EGTA. 20 TEACl. Intracellular tetraethylammonium chloride (TEA) was used to block K+ channels necessary for isolation the transduction current. To further isolate and increase the transduction current we also blocked voltage-gated sodium channels with 90 nM Tetrodotoxin (TTX) in the extracellular saline. The extracellular saline contained the following in mM: 185 NaCl, 10 KCl, 2 MgCl2, 2 CaCl2, 10 HEPES, 10 Trehalose, 10 Glucose. The saline was adjusted to pH 7.2 using NaOH. The osmolality of the intracellular and extracellular salines’ were 417 and 432 mOsm, respectively.

Whole-cell voltage-clamp recordings were performed with an EPC10-USB double patch-clamp amplifier (HEKA-Elektronik) controlled by the program Patchmaster (version 2 × 90.2, HEKA-Elektronik) running under Microsoft Windows (version 10). Electrophysiological data were sampled at 50 kHz. Voltage-clamp recordings were low-pass filtered at 2.9 kHz with a four-pole Bessel filter. Compensation of the offset potential were performed using the “automatic mode” of the EPC10 amplifier and the capacitive current was compensated manually. The calculated liquid junction potential between the intracellular and extracellular solutions was also compensated (15.6 mV; calculated with Patcher’s-PowerTools plug-in from www3.mpibpc.mpg.de/groups/neher/index.php?page=software). Series resistance was compensated at 77% with a time constant of 100 μs.

### Experimental design and statistical analysis

After hook electrode recordings the same locust was used for patch clamp recordings within 90 minutes. Throughout the manuscript n refers to the number of recorded neurons and N refers to the number of Müller’s Organ preparations used to achieve these recordings (i.e. n=10, N=6 means that 10 neurons were recorded from 6 Müller’s organs). All n numbers are displayed on the figures for clarity. The Spread of the data is indicated by 1 standard deviation as the standard deviation indicates the spread of the data, unlike standard error. Median and Q1 and Q3 are displayed by bars when individual measurements are plotted. For patch-clamp recordings, the treatment of the locust was blinded to the experimenter until after data analysis was complete. To test for differences and interactions between control, noise-exposed and aged locusts we used either a linear model (LM). The test statistic for these analyses (t) are reported with the p value. When the p value is below 0.05 the t and p value are highlighted in bold text.

## Results

### Injection of dsRNA knockdowns expression of nompC, nan and IFT88

We injected dsRNA complementary to *nompC, nan, ift88* and *gfp* into the locust’s abdomen both at 1- and 7-days post ecolsion (Figure 1A). We measured the relative change in gene expression of Müller’s organs. We found consistent and prolonged knockdown of *nompC, nan* and *ift88* compared to locusts injected with dsRNA against *Gfp* (Figure 1B). Both *nompC* and *nan* knockdown were decreased at 9 days post ecolsion when sound-evoked responses from the auditory nerve and transduction currents from individual auditory neurons were measured (Fig. 1Bi, ii). However, the knockdown of *nompC* was stronger compared to *nan* 48 hours after the second dsRNA injection (*nompC* t=7.2, nan t=4.5). Although the expression of *ift88*, was reduced after the first dsRNA injection it was not different 48 hours after the injection of dsRNA (Fig. 2Biii). We therefore also quantified expression of *ift88* 24 hours after the 2^nd^ injection when it was significantly reduced.

**Figure 1.**
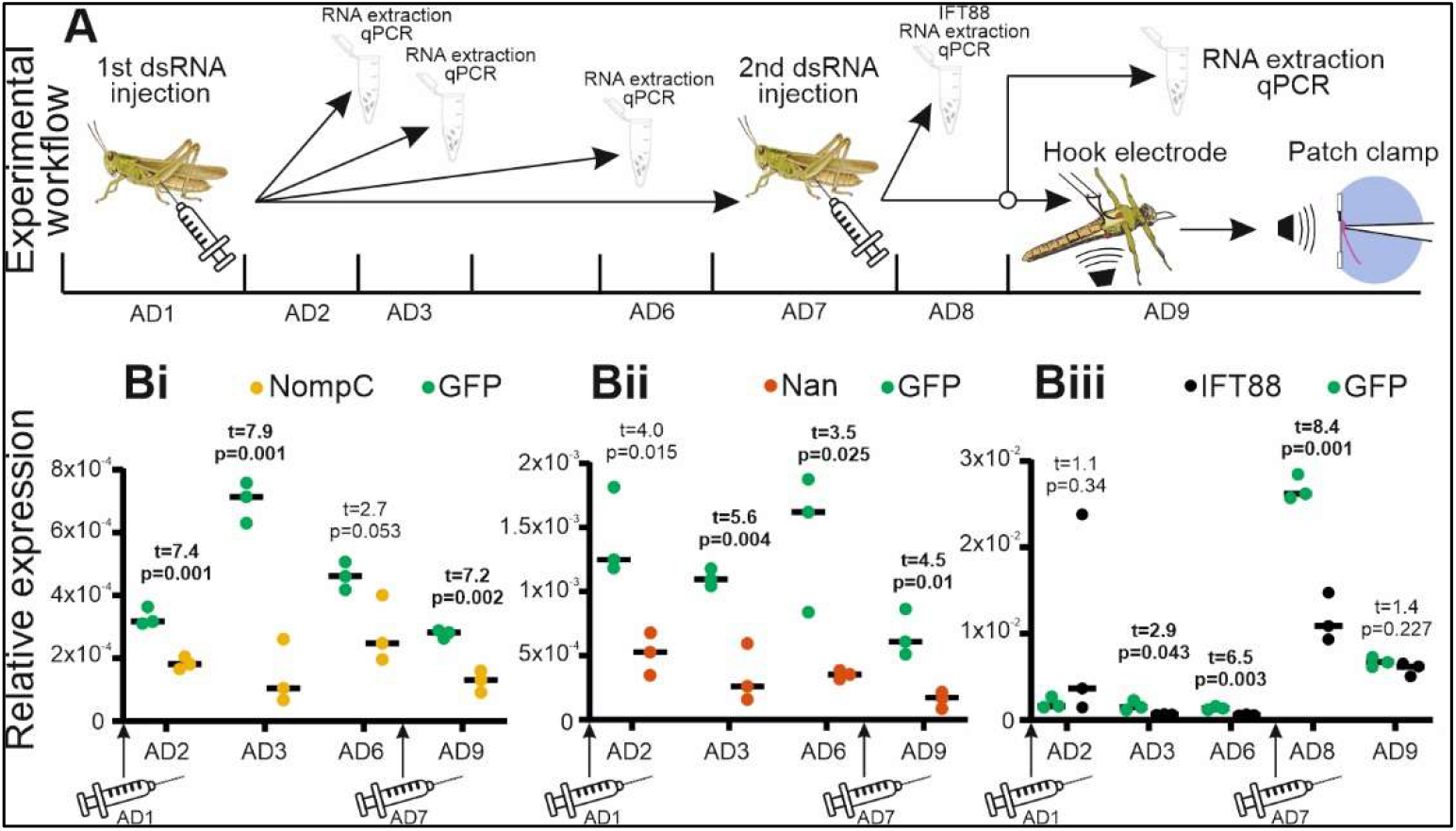
Experimental workflow and quantification of gene expression after knockdown. **A** Schematic of experimental workflow. N.B. after hook electrode recordings the same locusts were used for patch clamp recordings. **Bi** Relative expression after injection of dsRNA complementary to *nompC* and *gfp*, **Bii** Relative expression after injection of dsRNA complementary to *nan* and *gfp*, **Biii** Relative expression after injection of dsRNA complementary to *ift88* and *gfp*.

**Figure 2.**
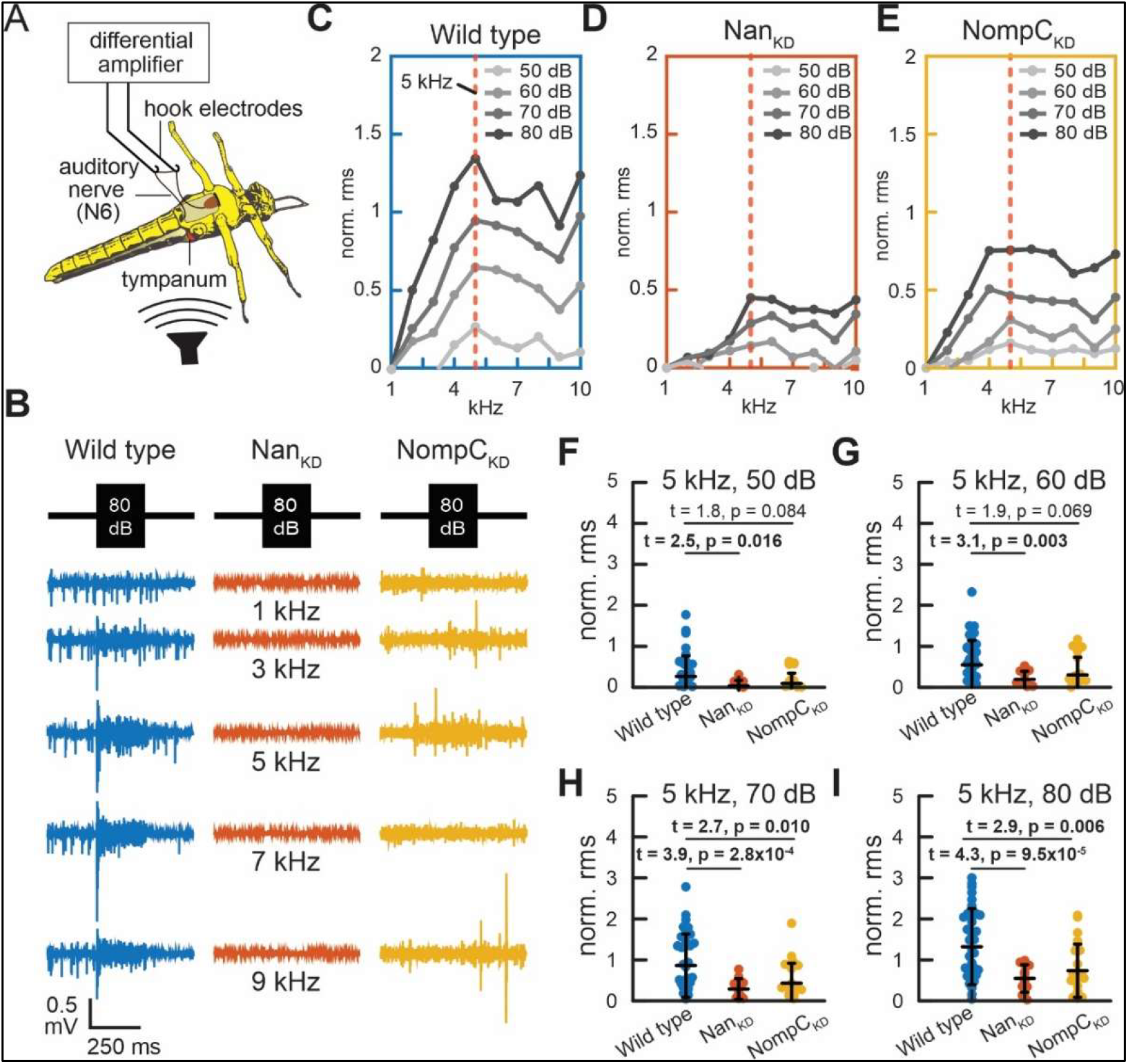
Hook electrode recordings from the auditory nerve upon pure-tone sound stimulation. **A** Schematic of a hook electrode recording from the auditory nerve N6 of the locust’s metathoracic ganglion. **B** Representative sound-evoked nerve responses at 80 dB SPL (Sound Pressure Level) from (left) wild type, (middle) *nan* knockdown *nan*_*KD*_, (right) and NompC-knockdown *nompC*_*KD*_ locusts. **C-E** Normalised Root-Mean-Square (RMS) of the sound-evoked response as a function of the stimulated frequencies (1-10 kHz) at increasing sound intensity (50-80 dB SPL) from the ensemble average of **C** wild type (N = 24, n = 37), **D** nan-knockdown *nan*_*KD*_ (N = 10, n = 12), and **E** NompC-knockdown *nompC*_*KD*_ locusts (N = 15, n = 23). The RMS nerve responses were normalised by mean RMS background activities of the auditory nerve before and after sound stimulation (see Materials and Methods). The red dashed line indicates 5 kHz, where the auditory nerve exhibits the largest nerve responses. **F-I** The corresponding scatter plots of the normalised RMS nerve responses at 5 kHz from **F** 50, **G** 60, **H** 70, and **I** 80 dB SPL. The superimposed horizontal lines indicate the ensemble mean (long line) and the standard deviation (short lines).

### Knockdown of nompC and nan resulted in decreased sound-evoked nerve potentials

We measured sound-evoked potentials from the auditory nerve of wild type locusts and locusts with either *nan* or *nompC* knocked down using hook electrode recordings (Fig. 2A). In response to pure tones the nerve produces adapting compound action potentials (Fig. 2B). We quantified the auditory nerve response by taking the root mean square normalised to the background nerve activity. We found that wild type locusts were most sensitive to 5 kHz (Fig. 2C), whereas *nan* and *nompC* mutants were more broadly sensitive to 5 kHz (Fig. 2D, E). The sound-evoked nerve response was quantified for SPLs between 50-80 dB for wild type, *nan* and *nompC* knockdowns (Fig. F-I). For the knockdown locusts we found decreased sound-evoked nerve response, compared to wild type controls, with the extent of the reduction increased for higher SPLs (T values = 2.9 and 4.3 at 80 dB SPL, T values = 1.8 and 2.5 at 50 dB SPL for *nompC* and *nan* respectively). The reduction in nerve activity was largest for *nan* knockdowns compared to *nompC* knockdowns across all SPLs.

### Knockdown of nompC or nan decreased transduction current but not discrete depolarisations

We measured electrophysiological properties, discrete depolarisations and ciliary-based sound-evoked transduction currents using an *ex vivo* preparation of the locust ear (Fig. 3A,B.) (Warren & Matheson, 2018). There were no differences in the resting membrane potential, membrane resistance or cell capacitance between wild type locusts, *nan* and *nompC* and *ift88* knockdown locusts (Table 1.) Intracellular recordings of chordotonal organ neurons are characterised by discrete depolarisations, small inwards currents of variable magnitude, assumed to be stochastic opening of the MET channel (Warren & Matheson, 2018). In response to pure tone stimulation auditory neurons of the similar desert locust produce an adapting transduction current (Warren & Matheson, 2018). We measured similar discrete depolarisations and an adapting sound-evoked transduction current in wild type migratory locusts in *nompC, nan* and *ift88*, knockdown migratory locusts (Fig. 3C). The transduction current was reduced in all three knockdown locusts compared to wild type locusts (Fig. 3Di). The discrete depolarisations were reduced in *ift88* knockdown locusts but the discrete depolarisations of *nan* and *nompC* knockdown locusts remained similar to wild type (Fig. 3Dii).

**Figure 3.**
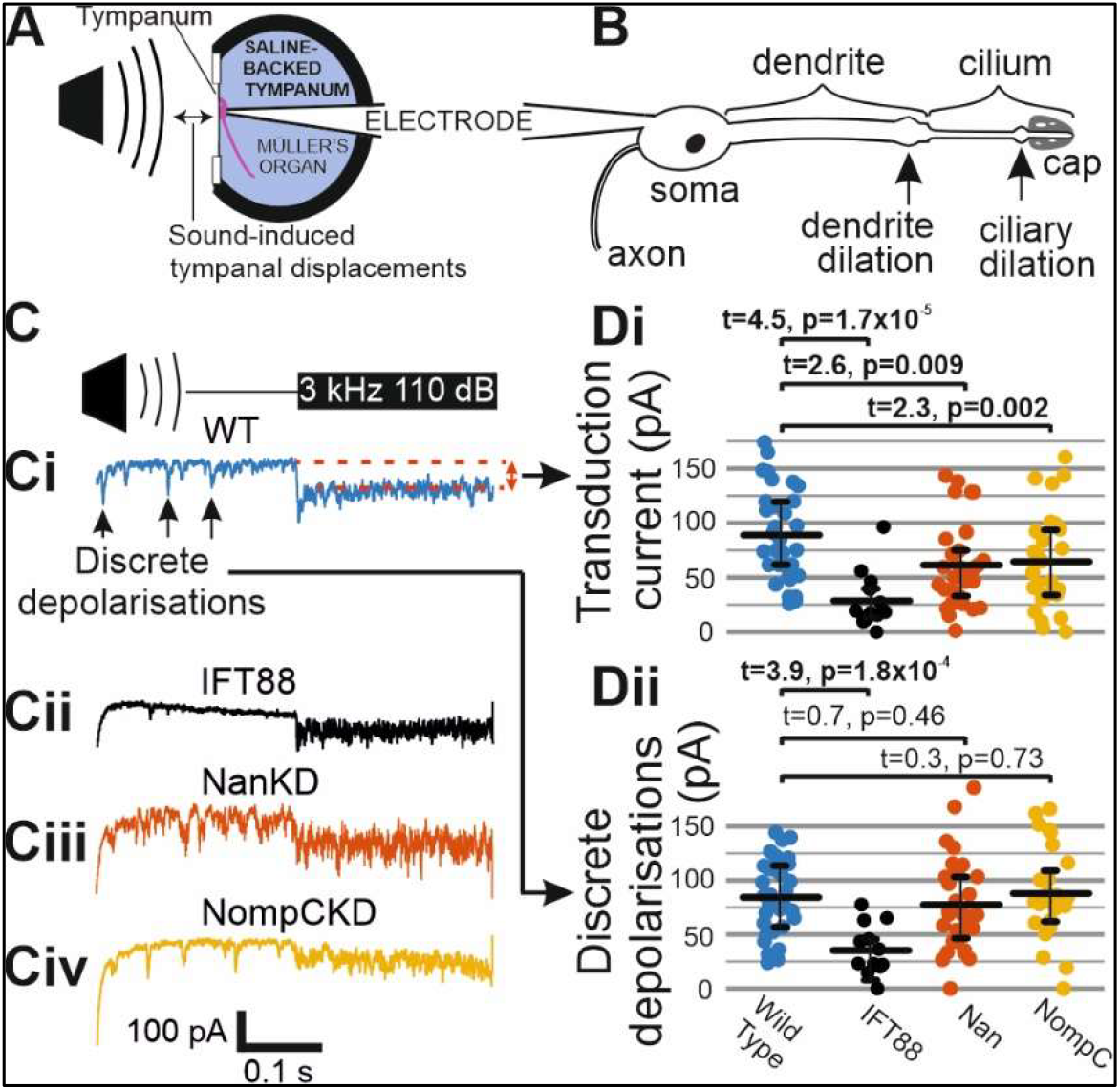
Patch-clamp recordings from auditory neurons that measure electrophysiological properties, discrete depolarisations and sound-evoked transduction current. A Schematic of *ex vivo* recording configuration, B Schematised morphology of auditory neuron. C Recordings of discrete depolarisations and sound-evoked transduction current at a holding potential of -100 mV for Ci wild type, Cii *ift88*, Ciii *nan* and Civ *NompC* dsRNA knockdown. Di Quantification of sound-evoked transduction current evoked with a 3 kHz 110 dB SPL tone and Dii discrete depolarisation (black lines represent median, Q1 and Q3, dof 98 and 97 respectively, Wild type N=12, n=23; *ift88* N=6, n=9; *nan*=9, n=29; *nompC* N=10, n=22).

**Table 1.** Electrophysiological properties of auditory neurons. There was no diference in electrophysiological properties in the auditory neurons of knockdown locusts compared to wild type.

| Treatment | Wild type | *ift88* | *nan* | *nompC* |
| --- | --- | --- | --- | --- |
| **Resting potential (mv)**<br>**dof=88** | **-46.3**<br>N=12, n=20 | **-42.7**<br>t=1.0, p=0.31<br>N=6, n=8 | **-42.9**<br>t=1.2, p0.21<br>N=9, n=29 | **-46.6**<br>t=0.12, p=0.91<br>N=10, n=19 |
| **Membrane resistance (MΩ)**<br>**dof=91** | **31.6**<br>N=12, n=20 | **26.5**<br>t=-1.2, p0.23<br>N=6, n=8 | **32.8**<br>t=0.35, 0.72<br>N=9, n=28 | **29.8**<br>t=0.54, p=0.59<br>N=10, n=21 |
| **Capacitance (pF)**<br>**dof=96** | **32.2**<br>N=12, n=23 | **35.9**<br>t=1.4, 0.16<br>N=9, n=8 | **31.0**<br>t=0.59, p0.56<br>N=9, n=29 | **33.8**<br>t=0.78, p=0.45<br>N=10, n=22 |

### Noise analysis of the transduction current does not conform to known MET channels

The variance (noise) in channel open probability should tend to zero when the channel is closed or constitutively open, whereas at half open probability the channel’s variance should be maximal (Qiu et al., 2025; Beurg et al., 2021). We measured discrete depolarisations and the transduction current in the desert locust as the amplitude of both is higher than the migratory locust. We found that variance is inherently high even without acoustic stimulation due to the discrete depolarisations even in the absence of acoustic stimulation (Figure 4A, B). Even when we use high intensity sound to maximise MET channel opening the variance plateaus and does not decrease further. This is in contrast to the MET channels of acoustic hair cells in mice where a parabolic function fits transduction current variance (Fig. 4B, red dotted line is parabolic function that extends to zero variance) (Beurg et al., 2021).

**Figure 4.**
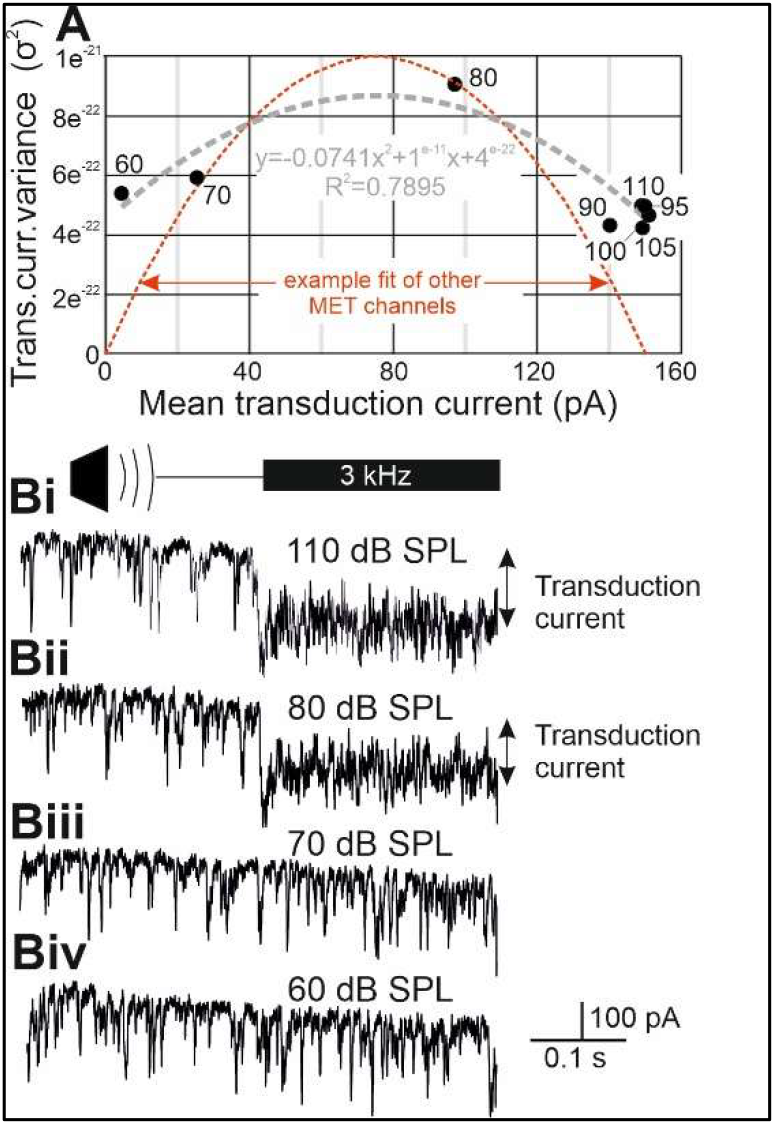
Noise analysis of transduction current. A. plot of mean transduction current vs variance. Numbers on graph donate the dB SPL of the tone to evoke the transduction current N.B. at 90 dB SPL and above the variance does not decrease further. Bi example stimulation at 110 dB SPL, Bii 80 dB SPL, Biii 70 dB SPL, Biv 60 dB SPL.

### Amplitude analysis of discrete depolarisations supports propagation towards the soma

We hypothesised that discrete depolarisations could be graded action potentials that travel along the cilium towards the soma. If this is true then the larger discrete depolarisations represent graded action potentials that have travelled further along the cilium towards the soma. We took advantage of the spatial limitation of voltage-clamp in the auditory neurons i.e. voltage-clamp was limited to close to the soma and the holding voltage (and driving force for depolarisation) decreased more distally along the dendrite and cilium. Due to this we predict that the magnitude of large discrete depolarisations would be increased proportionally more than the amplitude increase in smaller discrete depolarisations when the voltage was clamped from -60 to -100 mV. To test this we measured the magnitude of discrete depolarisations for 36 desert locusts and 83 auditory neurons for 3,234 voltage steps and analysed the distribution of their amplitudes on a histogram (Figure 5). The mean of the Gaussian distribution (µ) for the discrete depolarisations increased 18 pA from 56 to 74 pA at -60 mV and -100 mV but the larger infelction point increased proportionally more by 30 pA from 92 to 122 pA. This resulted in an increase in the skewness coefficient (α) of the Gaussian fit from 0.007 to 0.03 representing a disproportional increase for larger discrete depolarisations.

**Figure 5.**
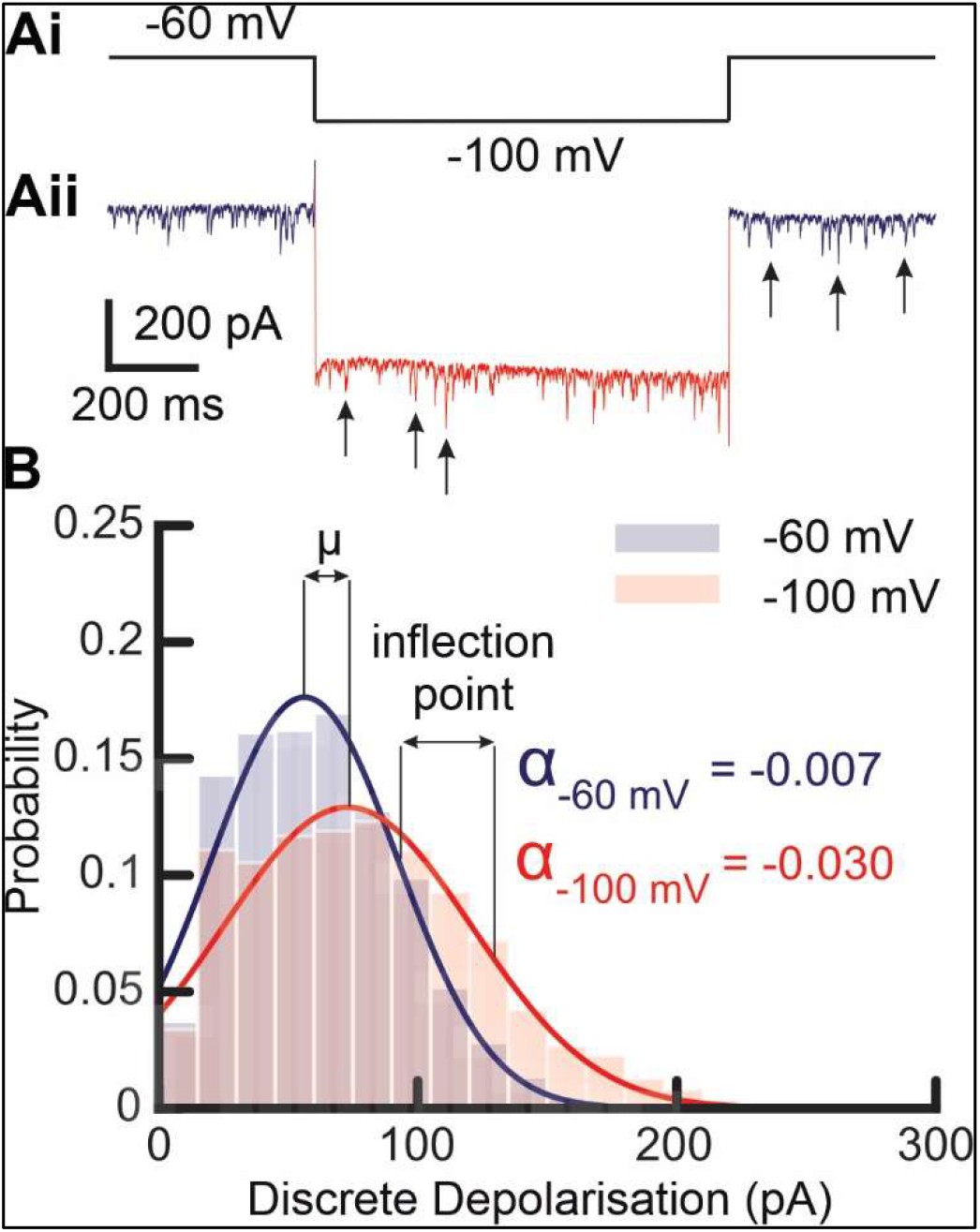
Amplitude of discrete depolarisations at an increased holding potential. Histogram plot and Gaussian fit of the amplitude of the discrete depolarisations at -60 mV (blue) and -100 mV (red).

## Discussion

We sought to discover the contribution of the ion channels NompC and Nan-Iav in auditory transduction the migratory locust. We knocked down the expression of *nan, nompC* and *ift88* and then recorded the sound-evoked nerve activity and the transduction current from individual auditory neurons in Müller’s organ of the migratory locust. We combined this with noise analysis from the highly similar auditory neurons of Müller’s organ of the desert locust.

### Discrete depolarisations

Previous intracellular recordings have assumed that the discrete depolarisations are elementary MET currents or stochastic openings of the transduction current (Hill, 1983; Warren & Matheson, 2018). Here, we show that knockdown of either *nompC* or *nan* did not affect the amplitude of discrete depolarisations casting doubts on the interpretation of patch-clamp recordings from chordotonal organ neurons (Warren & Matheson, 2018). As a positive control we knocked down *ift88*, a protein essential for transporting protein cargo along the cilium, which reduced discrete depolarisations and the transduction current. We therefore conclude that the discrete depolarisations are a third current carried by an unidentified ion channel.

We previously found all-or-nothing dendritic spikes and assume that the dendrite dilation is the spike initiation zone (Warren & Matheson, 2018). The dendritic spikes travel to the soma where they trigger the threshold for classical axonal spikes, presumably at the axon hillock. We hypothesise that the discrete depolarisations are ciliary based graded action potentials that are triggered in the ciliary dilation and travel along the basal cilium to trigger dendritic spikes at the dendrite dilation. Similar dilations termed “swellings” improve propagation of action potentials in Purkinje cells axons by decreasing spike failure rates (Lang-Ouellette et al., 2021), although modelling predicts a decrease in action potential velocity and peak height (Goldstein & Rall, 1974). Such a decrease in velocity at enlargements is modelled to cause an increase in spike “pile-up” that can cause the second spike to fail because it arrives during the refractory period of the first spike (Maia & Kutz, 2014). For graded potentials, however, a decrease in velocity would summate potentials in the dilation. We hypothesise that the ciliary dilation integrates MET currents which then triggers graded action potentials that propagate along the cilium to the dendrite dilation; the discrete depolarisations are similarly integrated at the dendrite dilation to trigger all-or-nothing dendritic spikes. Voltage-gated potassium and sodium channels, *shal* and *para* have recently been localised to the cilium in pupal Johnston’s organ in *Drosophila* (Gregory et al., 2025; Ravenscorft et al., 2023). The receptor lymph space of other insect mechanoreceptors is rich in potassium (Küppers, 1974) and it is assumed that the receptor lymph of chordotonal organ neurons is the same. Therefore, we predict that it is voltage-gated potassium channels that are responsible for the depolarisation phase of the discrete depolarisations.

### Noise analysis of the transduction current and amplitude changes in discrete depolarisations due to a voltage-step

We find it highly unlikely that the parabola fit to the noise from our sound-evoked currents predicts channel properties of the MET current. For instance, it predicts 5 channels with a conductance of ∼60 pA. This single channel current (60 pA) is at least three times higher than that measured for Nan-Iav or NompC but as the recording site is predicted to be many length constants distal this would make an unrealistic single channel conductance an order of magnitude higher ∼∼ 3000 pS, higher than any known eukaryotic ion channel. We suggest the reason that our noise analysis fails to give realistic values for the MET channel is that the sound-evoked currents, referred to as the transduction current, is composed of graded action potentials. This hypothesis is supported as large discrete depolarisations were proportionally increased compared to small discrete depolarisations when the electrical driving force was increased at the soma through a increase in the clamped voltage to more hyperpolarised potential of -100 mV. This is similar for voltage-clamped action currents (Hanson et al., 2004).

### The role of NompC and Nan-Iav

There are two running hypotheses of how auditory MET works in insect CO neurons, based on either NompC or Nan-Iav as the primary MET channel. The first hypothesis poises NompC as the primary MET channel and was derived from measurements of the mechanical fluctuations of the antennal Johnston’s organ in *Drosophila* but lacked electrical recordings (Gopfert et al., 2006). The counter hypothesis was derived from recordings of the summated currents from hundreds of auditory neurons and supports Nan-Iav but the recordings were electrotonically distant (Lehnert et al., 2013). Recordings directly from proprioceptive *Drosophila* larvae CO neurons combined with genetic knockout also support Nan-Iav as the primary MET channel (Li et al., 2021). Here, with patch-clamp recordings directly from the auditory neurons we find that both NompC and Nan-Iav contribute to the transduction current, with a slightly larger contribution from Nan-Iav. How can these disparate findings be reconciled? We suggest that *both* NompC and Nan-Iav form sound-sensitive ion channels. Two previous studies support two MET ion channels. The first predicted two MET ion channels in *Drosophila* Johnston’s organ, an insensitive and a sensitive population (Albert et al 2007). The second found that, under controlled stimulation of an individual ciliary ending, a bi-modal activation of the transduction current representing a sensitive and insensitive channel population (Chaiyasitdhi et al., 2025).

### Possible role of discrete depolarisations

Electrical potentials that travel along the cilium towards the soma were suggested 20 years ago but our results suggest they are not carried by Nan-Iav as hypothesised at the time (Göpfert et al., 2006). We suggest that discrete depolarisations are carried by voltage-gated potassium or calcium channels, the former recently localised to the cilium in *Drosophila* auditory neurons (Gregory et al., 2025). The receptor lymph that bathes the cilium is assumed to be high in potassium ions (opposite to conventional extracellular conditions). Therefore, the opening of potassium channels will lead to a depolarisation. There could also be a role of voltage-gated sodium channels, recently localised in the cilium of *Drosophila* larvae, to repolarise the cilium (Ravenscroft et al., 2023).

### Conclusion and predictions

We show that discrete depolarisations are not MET channel currents as knockdown of *nompC* or *nan* do not affect discrete depolarisation magnitude. Noise analysis of the discrete depolarisations suggest that they are graded action potentials that travel proximally to the soma. Both NompC and Nan-Iav are involved in generating the transduction current. We predict that the dilation found in the distal section of all Type 1 chordotonal organ neuron cilia is a charge integration centre.

